# Cholesterol accumulation-induced impairment of AKT signaling in LPS-stimulated macrophages play a dispensable role in suppressing HIF-1α-dependent glycolysis

**DOI:** 10.1101/2024.09.01.610717

**Authors:** Kenneth K.Y. Ting, Myron I. Cybulsky

**Author notes:** Correspondence; (1)-416-581-7483; PMCRT 3-306, 101 College Street, TMDT, Toronto, Ontario, Canada, M5G 1L7.

## Abstract

The formation of lipid-laden macrophages (Mφs) is a hallmark of atherosclerosis, yet how the accumulation of cholesterol in Mφs underlies the inflammatory process of atherogenesis remains unclear. It is well recognized that the reprogramming of metabolism in Mφs is critical for supporting their inflammatory responses, which may shed light on how the metabolism of Mφ foam cells is linked to inflammation. Indeed, recent research has now revealed Mφs that accumulate excess cholesterol adopt a distinct metabolic adaptation, a metabolic profile that is unexpectedly associated with a deactivated inflammatory response. Mechanistically, our group has previously shown that upon LPS stimulation, excess cholesterol accumulation in Mφs impaired their induction of AKT-dependent early glycolytic reprogramming and HIF-1α-dependent late glycolytic reprogramming. However, it remains unclear if these events are interconnected and synergistically contribute to the suppression of inflammation observed in these Mφs. Here, we demonstrated that cholesterol loading of Mφs impaired LPS-induced early glycolysis by reducing the phosphorylation of hexokinases, yet complete inhibition of AKT only modestly impaired HIF-1α-dependent glycolytic reprogramming. On the other hand, we confirmed that HIF-1α degradation, but not its reduced synthesis, is the primary mechanism that underlies its impaired expression in cholesterol loaded Mφs. Finally, we showed that cholesterol loading of Mφs alone was sufficient to induce oxidative stress, such as the production of 4-HNE, and deplete the levels of reduced KEAP1 proteins. Mφs lacking NRF2 resisted the effects of cholesterol loading on suppressing the expression of glycolytic and pro-inflammatory proteins.

## Introduction

Elevated plasma low density lipoprotein (LDL) is a key risk factor for atherosclerosis (1). LDL enters the artery intima, where both native and oxidized LDL (oxLDL) are taken up by macrophages (Mφs), resulting in foam cell formation (2). While the formation of foam cells in the artery intima is a hallmark of atherosclerosis, it is still debated whether foam cells or non-foamy Mφs are better at producing mediators that drive inflammation in atherosclerotic lesions. Traditional studies have shown that the accumulation of cholesterol within the Mφs alone is sufficient to induce inflammation (3), yet recent research findings have challenged this paradigm. For instance, our group has consistently shown in the past that cholesterol accumulation in Mφs not only fail to induce sterile inflammation, but it in fact impairs their ability to elicit a robust inflammatory response upon activation by an inflammatory stimulus, such as LPS (4-7). Notably, recent *in vivo* studies that have characterized the transcriptome of aortic intimal Mφs derived from atherosclerotic mouse models and human samples also confirmed that foamy Mφs were less inflammatory than non-foamy Mφs (8, 9). Yet, the mechanism behind how cholesterol loading suppresses the inflammatory responses in these Mφs is not well-understood.

In recent years, there is a growing appreciation that the rewiring of metabolic circuits in myeloid cells is a critical component to support their inflammatory response (10-12). For instance, many studies have now confirmed that both AKT-dependent early glycolysis and HIF-1α-dependent late glycolysis directly contribute to the inflammatory processes of myeloid cells by enhancing the production of inflammatory cytokines through transcriptional (10), translational (11) and epigenetic means (12). Given the intimate linkage between glycolysis and inflammation in Mφs, our group has also previously investigated how cholesterol loading of Mφ modulates their metabolic rewiring upon their activation by LPS and found that it impaired both AKT-dependent (5) and HIF-1α-dependent glycolysis (6). Specifically, we found the former was due to defective phosphorylation events upstream of AKT signaling (5), while the latter was due to the upregulation of NRF2-dependent antioxidative defense (6). However, it remains elusive how cholesterol loading-induced impairment of AKT signaling is linked to the suppression of early glycolysis. Moreover, AKT-mediated phosphorylation signaling through mTORC1 is known to regulate HIF-1α mRNA translation and its synthesis (13), thus it also remains unclear if the defective AKT signaling in cholesterol loaded Mφs can contribute to the suppression of HIF-1α-dependent glycolysis. Finally, it is also not well understood how cholesterol loading-induced oxidative stress activates the induction of NRF2-dependent antioxidative response.

In this study, we have now provided further evidence to support the concept that cholesterol loading of Mφs suppressed both AKT- and HIF-1α-dependent glycolytic reprogramming post LPS stimulation. Specifically, we found that cholesterol loading impaired LPS-induced early glucose uptake, AKT signaling and the phosphorylation of hexokinases. Yet, complete inhibition of AKT activity by pharmacological means only modestly inhibited HIF-1α-dependent-glycolytic reprogramming. On the other hand, cholesterol loading of Mφs impaired HIF-1α expression primarily through increasing its degradation, but not by decreasing its synthesis. Finally, we demonstrated that cholesterol loading alone induced oxidative stress and depleted the abundance of reduced KEAP1 proteins. Mφs that lack NRF2 were resistant to the suppressive effects of cholesterol loading on the expression of glycolytic and pro-inflammatory proteins.

## Methods

### Mouse strains

8 – 12 weeks-old mice were used. C57BL/6J (Strain #000664), B6.129S4(C)-*Vhl^tm1Jae^*/J (*Vhl*^fl/fl^) (Strain #012933), B6.129X1-*Nfe2l2^tm1Ywk^*/J (*Nfe2l2^-/-^*) (Strain #017009), B6.129P2-*Lyz2^tm1(cre)Ifo^*/J (Strain #004781) mice were purchased from The Jackson Laboratory. *Nfe2l2^-/-^* mice were generated by first crossing with wild type (WT) C57BL/6J mice, followed by heterozygote intercrossing. *Lys2-*Cre:*Vhl^fl/fl^* mice were generated by backcrossing a single *Lys2-*Cre transgene into *Vhl^fl/fl^* mice. Breeding for experiments consisted of crosses between Cre-positive and Cre-negative *Vhl^fl/fl^* mice. All mice were maintained in a pathogen-free, temperature-regulated environment with a 12-hour light and dark cycle and were fed a normal chow diet (NCD, 16 kcal% fat). All studies were performed under the approval of Animal User Protocols by the Animal Care Committee at the University Health Network according to the guidelines of the Canadian Council on Animal Care.

### Thioglycolate-elicited peritoneal Mφ (PMφ) isolation

Mice were injected intraperitoneally with 1 mL of 4% aged thioglycolate (ThermoFisher Cat#211716) and PMφs were harvested after 4 days by lavage with cold PBS containing 2% FBS. Cells were counted and cultured (37°C, 5% CO_2_) in DMEM supplemented with 10% FBS, 2 mM L-glutamine, 10,000 U/mL penicillin/streptomycin). Adherent PMφs were used in experiments after 18 h.

### Bone marrow derived Mφ (BMDMφ) generation

Mice were euthanized in a CO_2_ chamber and bone marrow cells were isolated from leg bones. Cells were cultured (37°C, 5% CO_2_) in RPMI supplemented with 10% FBS, 2 mM L-glutamine, 10,000 U/ml penicillin/streptomycin, and 40 ng/mL of M-CSF (PeproTech; Cat#AF-315-02) for 7 days. Cells were counted and replated for experiments.

### Transient transfection of RAW264.7 Mφs

RAW264.7 Mφs (2 x 10^6^, ATCC, Cat# ATCC TIB-71) were electroporated (Amaxa® Cell Line Nucleofector® Kit V, LONZA; Cat#VCA-1003) with 5 µg of HA-HIF1alpha (Addgene#18949) and HA-HIF1alpha P402A/P564A-pcDNA3 (Addgene#18955). Transfected cells were seeded in 6-well plates and cultured in recovery medium (DMEM, 20% FBS) for 3 h, then DMEM, 10% FBS for 48 h. G418 (400 µg/mL, replaced every 3 days, Gibco) was used for selection of stable-transfected cells After one week of selection, cells were used for experiments or cultured in DMEM,10% FBS, G418 (400 µg/mL).

### Lipid loading, LPS stimulation and inhibitor studies

PMφs were cultured for 24 h with cholesterol (50 μg/mL, Sigma Cat#C3045), followed by ultrapure LPS stimulation (10 ng/mL, InvivoGen, Cat#tlrl-3pelps) for up to 6 h. Ethanol (0.5%) was used as a carrier control for cholesterol. For inhibitor experiments, MK-2206 (0.5 µM) (Selleckchem, S1078), DM-NOFD (10µM) (Sigma-Aldrich, no. SML1874) and AKT2i (3 µM) (Sigma-Aldrich, no. 124029) was added 1 h prior to LPS stimulation. MG132 (10µM) (CST, #2194) was added at the 6^th^ hour of LPS stimulation, every 10 minute interval, to measure HIF-1α accumulation and hydroxylation.

### Immunoblotting

PMφs were cultured in each well of 12-well plates at 2 x 10^6^ per experimental condition. Cells were lysed in ice-cold RIPA buffer (1% NP40, 0.1% SDS, 0.5% deoxycholate in PBS, supplemented with 1 mM PMSF, 1X cOmplete™, EDTA-free Mini Protease Inhibitor Cocktail (Sigma Cat#11873580001) and 1X PhosSTOP™ (Sigma Cat#4906845001)) for 15 minutes. Protein concentrations in lysates were determined by Protein Assay Dye Reagent (BioRad Cat#5000006), diluted in 2x Laemmli sample buffer (BioRad Cat#161-0737) with fresh β-mercaptoethanol (BioRad Cat#1610710), and heated at 95°C for 5 minutes. Samples (20 μg of protein per lane) were resolved on 8%-15% SDS-PAGE gels and transferred to polyvinylidene difluoride membranes (Sigma Cat#IPVH00010) using a wet transfer system. Membranes were blocked with 5% skim milk non-fat powder or 3% BSA (Bioshop Cat#ALB003) in Tris-buffered saline-Tween (TBST) for 1 h at room temperature. Membranes were incubated with primary antibodies overnight: anti-Actin (Sigma, A2066), anti-GLUT1 (Abcam, ab115730), anti-Na,K-ATPase (CST#3010), anti-Phospho-Akt Substrate (RXXS*/T*) (CST#9614), anti-VDAC (CST#4866), anti-HIF-1α (CST#36169), anti-Hexokinase II (CST#2867), anti-Lamin A/C (CST#2032), anti-GAPDH (CST#5174), anti-NRF2 (CST#12721), anti-KEAP1 (ThermoFisher, #PA5-99434), anti-phospho-AKT Thr308 (CST#4056), anti-phospho-AKT Ser473 (CST#9271), anti-AKT (CST#9272), anti-phospho-ACLY Ser455 (CST #4331), anti-ACLY (CST #13390), anti-hydroxylated-HIF-1α (CST#3434), anti-TXNIP (ThermoFisher, MA532771), anti-4-HNE (ThermoFisher, MA5-27570), anti-Nox2 (Abcam, 310337), followed by washing and incubation with HRP-conjugated anti-rabbit IgG (CST#7074) (22°C, 1 h). Blots were developed using Immobilon Forte Western HRP substrate (Sigma, WBLUF0100), imaged with Microchemi 4.2 (BioRad) and analyzed with ImageJ. To assess GLUT1 and phospho-hexokinase expression in different subcellular compartments, subcellular fractionation followed by immunoblotting was performed (14). To assess oxidized and reduced KEAP1 protein expression, redox western blotting was performed (15).

### Extracellular Acidification Rate and Oxygen Consumption Rate (OCR) measurement

PMφs (2.5 x 10^5^) were cultured in XF24 well plates (Agilent Technologies, Cat#102342-100), incubated with cholesterol for 24h or MK2206 for 1h, followed by the injection of LPS by the Seahorse analyzer. Real-time ECAR and OCR values were acquired and normalized (baselined OCR %) to one reading before **RNA isolation and real-time (RT) PCR.** Total RNA was isolated with E.Z.N.A.® Total RNA Kit I (Omega Cat#R6834-01) and reverse transcription (RT) reactions were performed with High-Capacity cDNA Reverse Transcription Kit (ThermoFisher Cat#4368814) according to manufacturer’s protocol. Real time quantitative PCR (qPCR) was then performed using a Roche LightCycler 480 with Luna® Universal qPCR Master Mix (New England Biolabs, Cat#M3003E). Quantification of mRNA was performed by using primers that span over two adjacent exons, quantified using the comparative standard curve method and normalized to hypoxanthine phosphoribosyltransferase (HPRT), a housekeeping gene. Primer sequences used for qPCR are listed below:

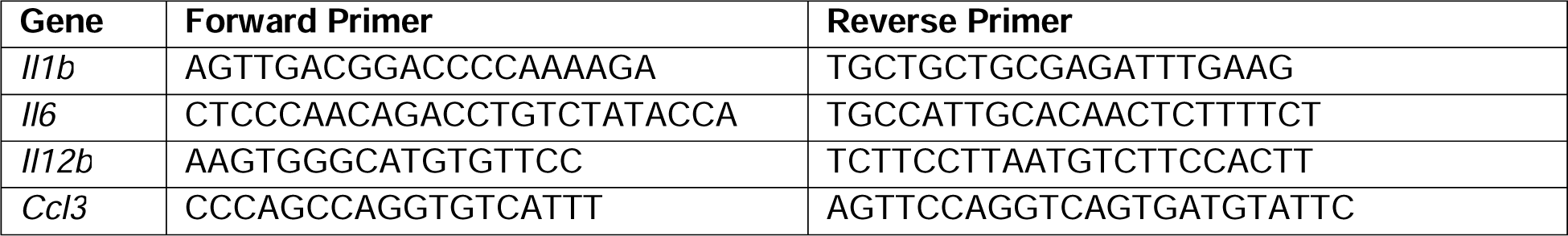
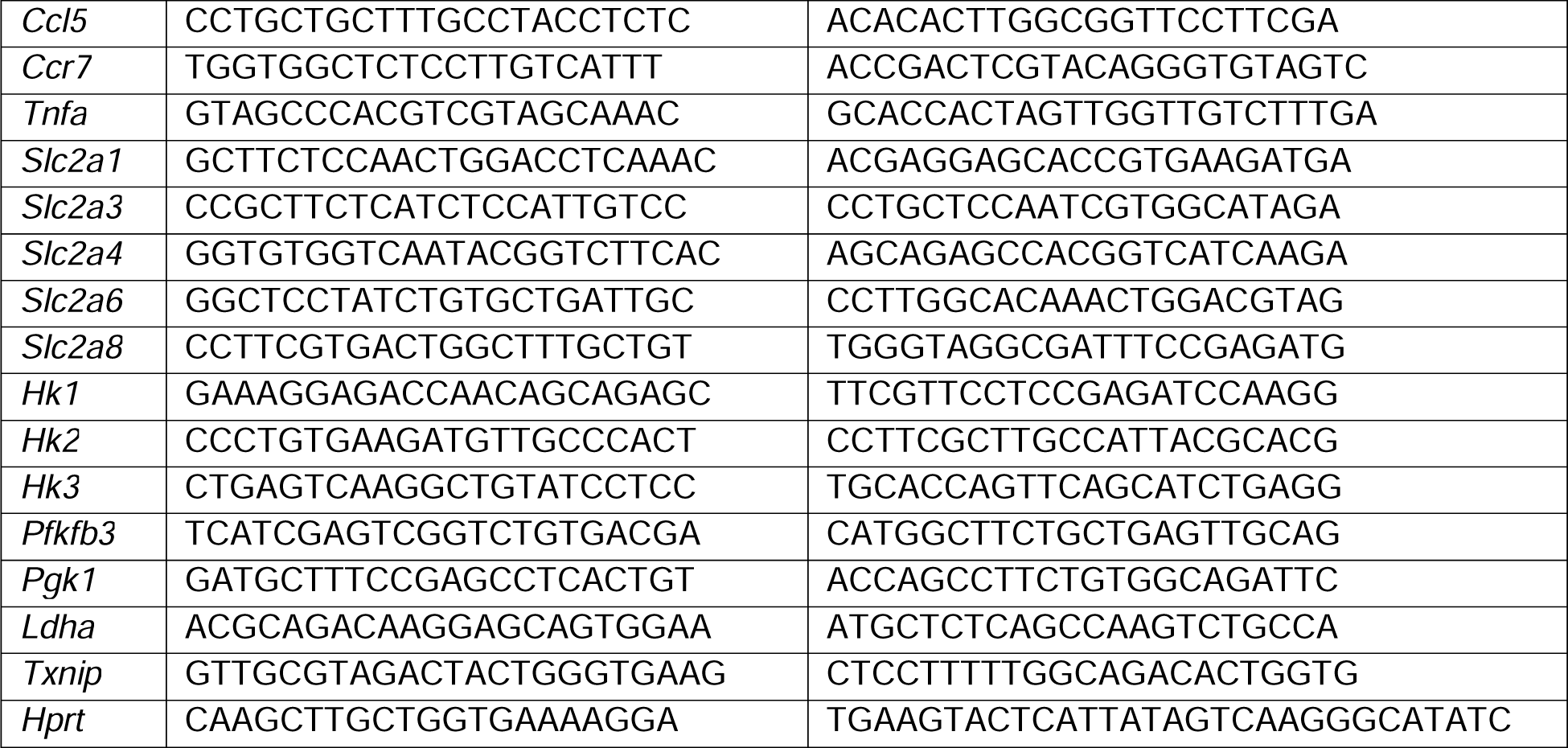

### Glucose uptake and immunofluorescence microscopy

PMφs (3 x 10^6^) were first seeded in 35 mm petri dish, with 14 mm microwell (MatTek, P35G-1.5-14-C), then cultured with cholesterol overnight. Cells were stimulated with LPS for 3 h the next day, washed three times with pre-warmed HBSS (Wisent, Cat#311-513-CL). To detect glucose uptake, cells were cultured for 1 h at 37°C, 5% CO_2_in DMEM without glucose (ThermoFisher #11966025) supplemented with 10% FBS, 2-NBDG (Cayman Chemicals #11046) (100 μg/mL) and 32.4 μM of Hoechst nuclear staining reagent (ThermoFisher #H3570). Cells were imaged with Olympus FluoView 1000 Laser Scanning Confocal Microscope (Olympus America) or A1R Confocal microscope with resonant scanner (Nikon). Mean fluorescence intensity measurements represented the ratio of total fluorescence intensity for each field to the number of nuclei in that field.

### Statistical analysis

The statistical test(s) used in each experiment is listed in the figure legends. In general, the figures show pooled data from independent experiments. All experiments were repeated at least three times and the number of biological replicates is indicated as the n value. Statistical analyses were performed using the Prism software, unless otherwise specified in the figure legends.

## Results

### Cholesterol loading of PMφs impaired LPS-induced AKT-dependent early glycolytic reprogramming

Our past study has shown that cholesterol loading of thioglycollate-elicited peritoneal Mφs (PMφs) impaired AKT-dependent early glycolytic reprogramming post LPS stimulation (5), yet how the impairment of AKT signaling is linked to glycolysis remains unclear. To elucidate the mechanism, we first confirmed that cholesterol loading of PMφs suppressed the induction of glycolysis post LPS stimulation by utilizing the seahorse analyzer. As shown in Figure 1A and Figure 1B, the analyzer revealed that LPS stimulation of PMφs significantly increased the extracellular acidification rate (ECAR), a surrogate measure of the glycolysis, and decreased oxygen consumption rate (OCR), a surrogate measure of mitochondrial respiration, in a time-dependent manner. This is consistent with the past literature findings that demonstrated the Warburg effect in LPS-activated Mφs, where these cells engage a metabolic switch from mitochondrial respiration to aerobic glycolysis to fuel their energetic demands (16). However, PMφs that were previously loaded with cholesterol, their induction of glycolysis was impaired post LPS stimulation, while their mitochondrial respiration activity only was modestly inhibited. Taken together, these data suggest that cholesterol loading blocked the metabolic shift of LPS-activated Mφs from mitochondrial respiration to glycolysis. To elucidate the mechanism, we first focused on the molecular events that take place during the early phase of glycolytic reprogramming (0-4h post LPS stimulation), such as glucose uptake (17). To measure glucose uptake, we incubated PMφs with 2-(7-Nitro-2,1,3-benzoxadiazol-4-yl)-D-glucosamine (2-NBDG), a fluorescent glucose analog, in glucose-free media and performed immunofluorescence microscopy to quantify its intracellular levels post LPS stimulation. As shown in Figure 1C, LPS stimulation significantly induced the intracellular uptake of 2-NBDG in PMφs, yet this induction was significantly reduced in cholesterol loaded PMφs. Interestingly, the uptake of 2-NBDG was also reduced in cholesterol loaded PMφs prior to LPS stimulation. Understanding that the steady-state abundance of intracellular glucose is determined by both its uptake and intracellular retention, which are regulated by glucose transporters and hexokinases respectively, we explored if cholesterol loading affected the expression and activity of these glycolytic proteins. For glucose transporters, we first performed qPCR to assess the gene expression of various members of the glucose transporter family and found that LPS stimulation induced the expression of *Slc2a1*(GLUT1) and *Slc2a6* (GLUT6), in which their expression was suppressed by cholesterol loading (Figure 1D). We then chose to investigate GLUT1 instead of GLUT6 as others have already shown that GLUT6 is localized to lysosome membranes and does not play a role in mediating glucose uptake (18). To investigate if cholesterol loading modulated GLUT1 expression on the plasma membrane, we performed subcellular fractionation to isolate cytosolic, plasma membrane and total membrane components, followed by immunoblotting of GLUT1. As shown in Figure 1E, the expression of GLUT1 in all fractions was not affected by LPS nor by cholesterol loading, thus suggesting the unlikelihood that cholesterol loading of PMφs suppresses LPS-induced glucose uptake via GLUT1. We next turned our attention to hexokinases as its phosphorylation of glucose is critical to mediate its entrapment, and thus can modulate intracellular glucose levels. Notably, the phosphorylation levels of hexokinases, which is an indicator of the activity of hexokinases (19), has been previously demonstrated to be increased in LPS-activated dendritic cells (11). To explore if cholesterol loading affected the phosphorylation of hexokinases, we performed subcellular fractionation to isolate cytosolic and mitochondrial components, followed by immunoblotting of phosphorylation of hexokinases. As shown in Figure 1F, LPS stimulation increased the phosphorylation levels of hexokinases in the mitochondrial fraction, but not in the cytosolic fraction, and this enhancement was impaired in cholesterol loaded PMφs. Since AKT has been shown to phosphorylate hexokinases and stimulate their activity (19), we measured its activation, which is reflected by the phosphorylated levels on its Thr308 residue, and confirmed that its induction by LPS was suppressed by cholesterol loading (Figure 1G). The activity of AKT, which can be measured by the phosphorylated levels of Ser455 on ACLY, a direct downstream target of AKT (20), was also impaired by cholesterol loading post LPS stimulation. Overall, our results have demonstrated that cholesterol loading of PMφs impaired LPS-induced glucose uptake, AKT activation and hexokinases activity.

**Figure 1.**
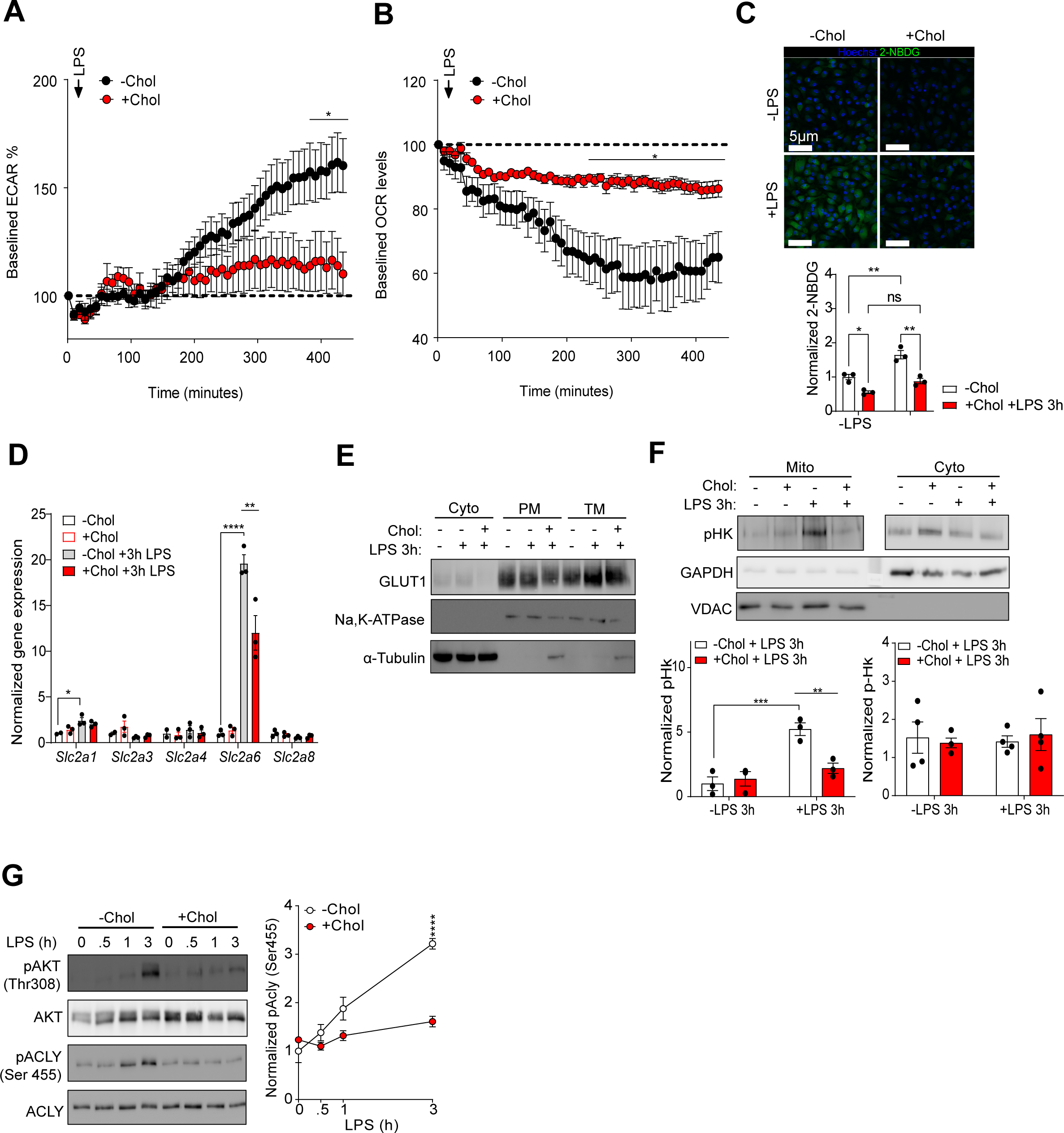
Cholesterol loading of PMφs impaired LPS-induced AKT-dependent early glycolytic reprogramming. **A and B)** Seahorse analyzer provides real-time measurements of baselined ECAR (A) and OCR (B) measurements in ±Chol loaded PMφs during 6h of LPS stimulation (n=6-7). Arrow indicates the time of LPS injection. Dashed line indicates the baselined value. **(C)** Immunofluorescence microscopy that illustrates 2-NBDG (Green) intracellular accumulation and nuclei staining by Hoechst 33342 in ±Chol loaded PMφs post 3h of LPS stimulation (n=3). **(D)** qPCR analysis that shows the gene expression of various glucose transporters in ±Chol loaded PMφ post 3h of LPS stimulation (n=3). **(E)** Immunoblot analysis that shows the expression of GLUT1 in ±Chol loaded PMφs post 3h of LPS stimulation in cytoplasmic (Ctyo), plasma membrane (PM) and total membrane (TM) fractions (n=2). Na,K-ATPase was used as a loading control for membrane fractions. α-tubulin was used as a loading control for cytosolic fraction **(F)** Immunoblotting analysis that shows the expression of phospho-hexokinase in ±Chol loaded PMφs ±LPS 3h stimulation in mitochondrial and cytoplasmic fraction (n=3). GAPDH and VDAC were used as a loading control for cytoplasmic and mitochondrial fraction respectively. **(G)** Immunoblotting analysis that shows the expression of pAKT (Thr308), AKT, pACLY (Ser455) and ACLY in ±Chol loaded PMφs within 3h of LPS stimulation (n=4). The mean ± SEM is plotted in all graphs. Significant differences were determined by two-way ANOVA with Bonferroni post hoc test in all graphs (* P < 0.05, ** P < 0.01, *** P < 0.001, **** P < 0.0001).

### Cholesterol loading of PM**φ**s impaired LPS-induced HIF-1**α**-dependent late glycolytic **reprogramming independent of the defective AKT signaling**

Past research has shown that LPS-induced Mφs adopt a two-phase transition of glycolytic reprogramming (17), where the early phase is rapid and regulated by AKT-mediated phosphorylation events (11), while the late phase is more robust and regulated by HIF-1α-mediated transcription of glycolysis genes (10).

However, it remains unclear if a defective AKT signaling cascade can hamper the induction of subsequent HIF-1α-mediated glycolytic reprogramming. Since past research has shown that the 5’ untranslated region of HIF-1α mRNA has an oligopyrimidine (TOP) motif that enables its translation to be regulated by mammalian target of rapamycin complex 1 (mTORC1) (13), which is downstream of AKT signaling, it raises the possibility that the defective AKT signaling in cholesterol loaded PMφs may suppress subsequent HIF-1α-dependent glycolytic reprogramming. To determine this, we measured HIF-1α protein levels in LPS-stimulated PMφs upon complete functional blockade of AKT function. As shown in Figure 2A, complete blockade of AKT function with pan-AKT inhibitor, MK2206, only modestly impaired LPS-induced HIF-1α levels. To assess the effects of AKT inhibition on HIF-1α-dependent glycolytic reprogramming, we also measured ECAR and OCR in LPS-stimulated Mφs treated with MK2206. As shown in Figure 2B, basal and LPS-induced ECAR was only modestly suppressed despite complete blockade of AKT activation by MK2206. With regards to the expression of pro-inflammatory genes, such as *Il1b*, which is transcriptionally regulated by HIF-1α (10), was only partially inhibited by MK2206 (Figure 2C). Since we have also previously shown that cholesterol loading of Mφs selectively impaired the activity of AKT2, one of the three isoforms, post LPS stimulation (5), we also explored the possibility that selective AKT2 signaling may contribute suppressing HIF-1α-dependent glycolysis. As shown in Supplementary Figure 1A, while the inhibitory effects of AKT2 on LPS-induced inflammatory gene expression was more potent than MK2206, it did not affect the expression of many HIF-1α-targeted glycolysis genes (Supplementary Figure 1B). This complemented with our previous findings that selective inhibition of AKT2 did not affect LPS-induced glycolysis (5). To further assess the potential contributory role that AKT may play in suppressing the synthesis of HIF-1α protein levels, we measured the synthesis rate of HIF-1α protein levels by blocking its proteasomal degradation with MG132 in a time-dependent manner. As shown in Figure 2D, the rate of HIF-1α protein accumulation is comparable between control and cholesterol loaded PMφs post LPS stimulation. On the other hand, the rate of HIF-1α hydroxylation on Pro577, which is catalyzed by prolyl hydroxylases and mediates HIF-1α degradation, is much higher in cholesterol loaded PMφs. Therefore, these results collectively demonstrate that cholesterol loading-induced impairment of AKT signaling is unlikely to play a major role in modulating subsequent HIF-1α-dependent glycolysis. To verify that the enhancement of HIF-1α degradation is the primary mechanism that led to its impairment, we stably transfected two plasmids separately into cholesterol-loaded RAW264.7 Mφ cell lines, one that encodes for WT HIF-1α proteins, and the other one encodes for a non-degradable HIF-1α protein mutant. As shown in Figure 2E, immunoblot analysis of HIF-1α protein levels showed that cholesterol loading significantly inhibited the cell line that expressed the levels of WT HIF-1α proteins, but did not suppress the cell line that expressed the levels of non-degradable HIF-1α mutant proteins. Finally, to assess the role of HIF-1α in regulating LPS-induced inflammatory and glycolysis gene expression, we have utilized *Lyz2Cre:Vhl^fl/fl^* mice to acquire Mφs that lack VHL, an adaptor protein that mediates HIF-1α proteasomal degradation. Our previous study has already confirmed that Mφs that lack VHL were resistant to the suppressive effects of cholesterol loading on HIF-1α protein expression, yet their expression of pro-inflammatory and glycolytic genes were still suppressed (6). We later demonstrated that cholesterol loading induced the activity of Factor Inhibiting HIF (FIH), which impaired the transactivation capacity of HIF-1α, and that the inhibition of FIH was also required to restore HIF-1α transcription responses, in addition to blocking HIF-1α degradation (6). However, this was only demonstrated in RAW264.7 Mφ cell line and it remains unclear if similar findings would also be observed in primary Mφs. To validate this, we again utilized primary Mφs that lack VHL and pharmacologically inhibited FIH with an inhibitor (FIHi), known as DM-NOFD, and confirmed that expression of selected pro-inflammatory and glycolysis genes were reinstated upon FIH inhibition (Figure 2F). Taken together, these data collectively demonstrate that while cholesterol loading of PMφs impaired LPS-induced AKT signaling, it is unlikely to contribute to the subsequent suppression of HIF-1α transcriptional responses.

**Figure 2.**
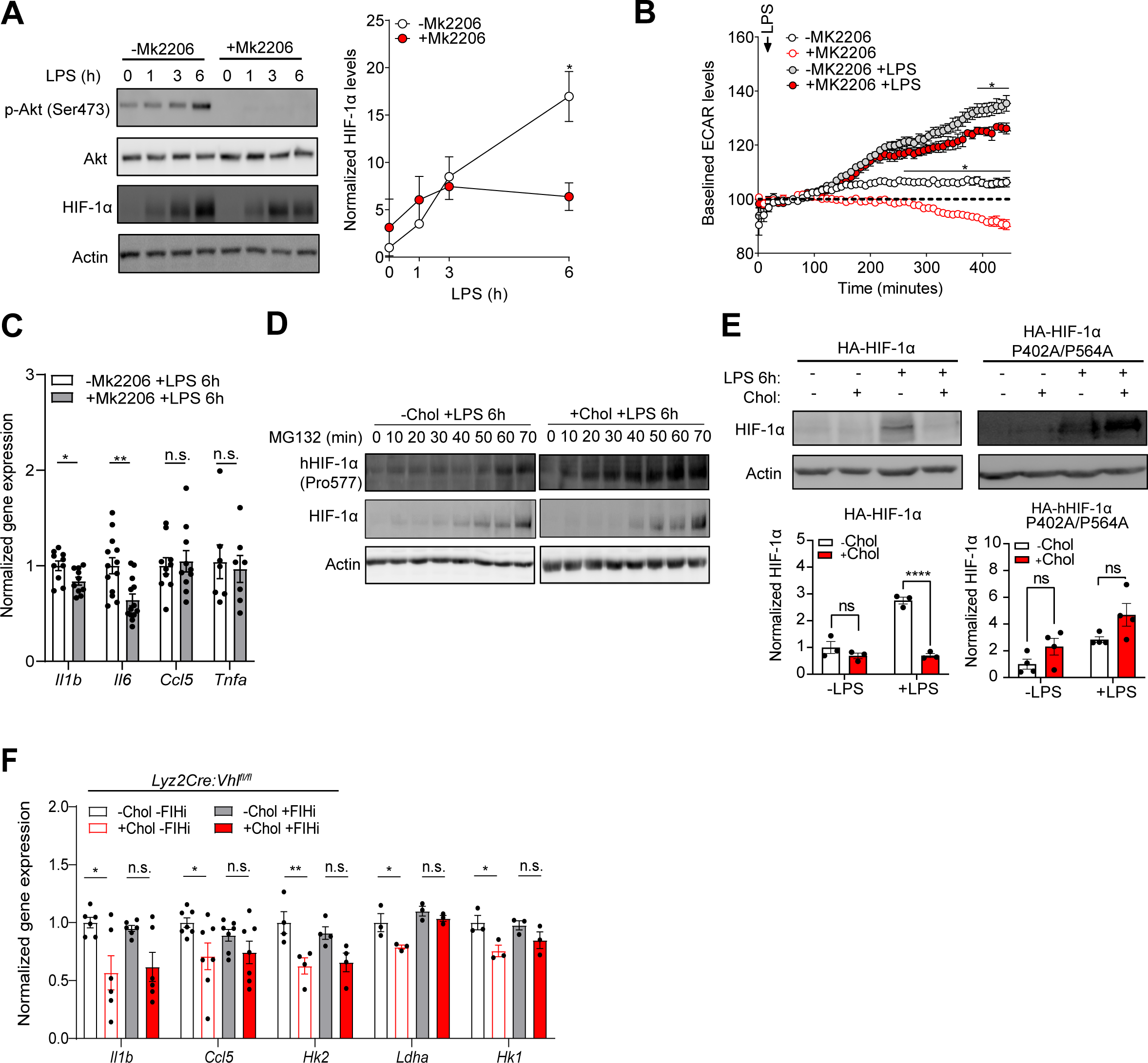
Cholesterol loading of PMφs impaired LPS-induced HIF-1α-dependent late glycolytic reprogramming independent of the defective AKT signaling. **(A)** Immunoblotting analysis that shows pAKT (Ser473), AKT and HIF-1α protein levels in ±MK2206 treated PMφs within 6h of LPS stimulation (n=2). MK2206 was added 1h before LPS stimulation **(B)** Seahorse analyzer shows the baselined ECAR levels in ±MK2206 treated and ±LPS stimulated PMφs (n=4). **(C)** qPCR analysis that shows the gene expression of *Il1b, Il6, Ccl5* and *Tnfa* expression in ±MK2206 treated PMφs post 6h of LPS stimulation (n=6-10). MK2206 was added 1h before LPS stimulation. **(D)** Immunoblot analysis of hydroxylated HIF-1α and HIF-1α protein levels in ±Chol loaded PMφs post 6h of LPS stimulation (n=4). MG132 was added on the 6^th^ hour of LPS stimulation at every 10-minute intervals. **(E)** Immunoblot analysis of HIF-1α protein levels in ±Chol loaded ±LPS 6h treated RAW264.7 Mφ cell lines that stably express plasmids encoding HA-HIF-1α proteins and HA-HIF-1α P402A/P564A proteins (n=4). **(F)** qPCR analysis that shows the gene expression of inflammatory and glycolysis genes in ±Chol loaded ±FIHi treated BMDMφs from *Lys2Cre:Vhl^fl/fl^*mice post 6h of LPS stimulation. FIHi was added 1h prior to LPS stimulation (n=3-6). The mean ± SEM is plotted in all graphs. Significant differences were determined by two-way ANOVA with Bonferroni post hoc test in **(A, B, E)**, and by unpaired Student’s *t*-test in **(C and F)** (* P < 0.05, ** P < 0.01, **** P < 0.0001).

### Cholesterol loading of PM**φ**s activated oxidative stress and depletion of reduced KEAP1 proteins prior to LPS stimulation

Our past research has demonstrated that cholesterol-loading induced NRF2-dependent antioxidative response underlies the impairment of HIF-1α-dependent response (6). More importantly, we have further demonstrated that the cysteine residues of KEAP1, a negative regulator of NRF2 that mediates its proteasomal degradation, were dysfunctional in cholesterol loaded Mφs and this contributed to the enhancement of NRF2-dependent response (6). However, it remains unclear how cholesterol loading-induced oxidative stress led to the dysfunction of cysteine residues and the molecular events that consequently take place. To explore this, we first confirmed that cholesterol loading of PMφs alone is sufficient to induce oxidative stress. As shown in Figure 3A, seahorse analyzer demonstrated that 12h of cholesterol loading of PMφs induced a higher mitochondrial respiration activity than treating Mφs with ethanol carrier. This finding complemented with our past results that showed cholesterol loading of PMφs induced the most mitochondrial-derived ROS levels at 12h post of lipid loading (6). Notably, we measured the protein expression of Thioredoxin interacting protein (TXNIP), a protein that indicates the presence of oxidative stress and is a transcriptional target of NRF2 (21), and found its levels were induced by cholesterol loading alone and sustained post LPS stimulation (Figure 3B). Collectively, these data confirm with our previous findings that cholesterol loading of PMφs alone induced NRF2-dependent oxidative stress response prior to LPS stimulation. To determine how cholesterol loading-induced oxidative stress is linked to the dysfunction of KEAP1 cysteine residues, we explored the role of lipid peroxidation products in interfering with KEAP1 cysteine residues sensory functions. Among all, we focused on 4-HNE as its role in inhibiting KEAP1 cysteine residues is well-established (22, 23). As shown in Figure 3C, cholesterol loading of PMφs alone induced a significantly higher level of 4-HNE modified proteins, thus suggesting the possibility that 4-HNE production was increased in cholesterol loaded PMφs. To further examine how KEAP1 function is affected by cholesterol loading, we performed redox western blotting to assess the relative levels of oxidized and reduced KEAP1 levels, in which the oxidized KEAP1 proteins enable NRF2 to be stabilized while the reduced KEAP1 proteins mediate NRF2 to undergo proteasomal degradation. As shown in Figure 3D, we first optimized our redox western blotting assay with H_2_O_2_, where its addition rapidly induced oxidative stress in PMφs, as detected by an initial rise of oxidized KEAP1 proteins and a decline of reduced KEAP1 proteins. As the antioxidative defense response is activated and redox homeostasis is later restored, the induction of oxidized KEAP1 proteins also subsided, followed by a restoration of reduced KEAP1 protein levels. Utilizing this redox western blotting assay, we then characterized the relative levels of oxidized and reduced KEAP1 proteins in cholesterol loaded PMφs. As shown in Figure 3E, during the 24h course of cholesterol loading, we detected a significant rise of oxidized KEAP1 protein levels and a corresponding decrease of reduced KEAP1 protein levels. Notably, almost all KEAP1 proteins were oxidized by the end of the cholesterol loading kinetic. This is consistent with our past observation that NRF2 levels were induced by cholesterol loading alone (5). Finally, to repeatedly demonstrate the role of NRF2 in suppressing HIF-1α-dependent glycolysis and inflammation, we utilized Mφs that lack NRF2 and confirmed that these cells were resistant to the inhibitory effects of cholesterol loading on the protein expression of Hexokinase II and NAPDH-Oxidase 2, which are known transcriptional targets of HIF-1α.

**Figure 3.**
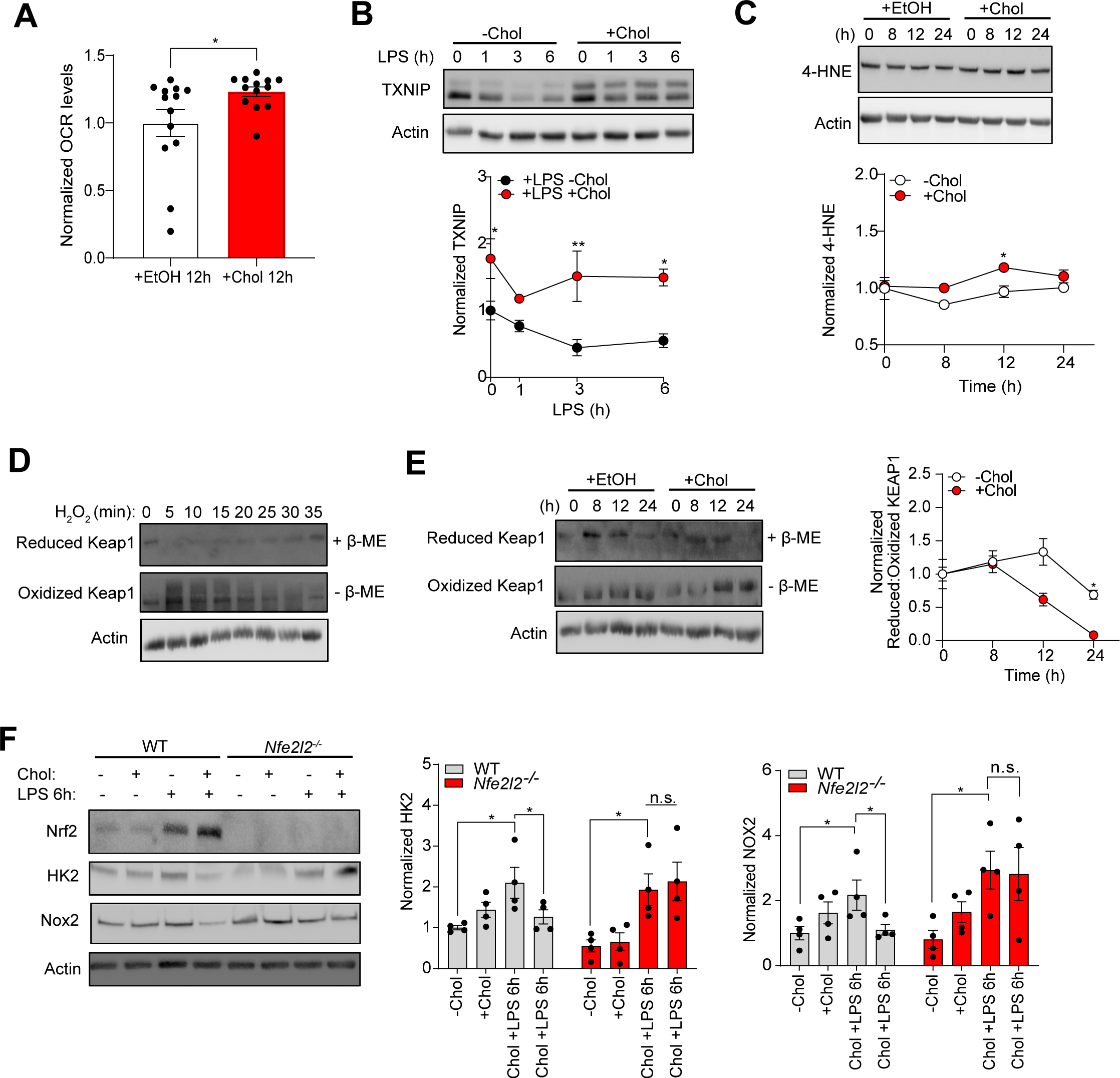
Cholesterol loading of PMφs activated oxidative stress and depletion of reduced KEAP1 proteins prior to LPS stimulation. **(A)** Seahorse analysis shows OCR measurements in ±Chol loaded PMφs post 12h of lipid loading (n=13). **(B)** Immunoblotting analysis of TXNIP protein levels in ±Chol loaded PMφs post 6h of LPS stimulation (n=4). **(C)** Immunoblotting analysis of 4-HNE adducted protein levels in ±Chol loaded PMφs within 24h of lipid loading (n=3). **(D)** Redox western blotting analysis of reduced and oxidized KEAP1 protein levels in PMφs treated with H_2_O_2_ in a time-dependent manner. The reduced KEAP1 proteins were detected under reducing conditions (with β-ME) while oxidized KEAP1 proteins were detected under non-reducing conditions (without β-ME). **(E)** Redox western blotting analysis of reduced and oxidized KEAP1 protein levels in ±Chol loaded PMφs within 24h of lipid loading. **(F)** Immunoblot analysis of NRF2, HK2 and NOX2 protein levels in WT and *Nfe2l2^-/-^* PMφs ±Chol loading ±6h of LPS stimulation (n=4). The mean ± SEM is plotted in all graphs. Significant differences were determined by two-way ANOVA with Bonferroni post hoc test in **(B, C, E and F)**, and by unpaired Student’s *t*-test in **(A)** (* P < 0.05, ** P < 0.01).

## Discussion

Mφs, a group of leukocytes that constitutes our innate immunity, are also major contributors of atherosclerosis. Traditional studies have shown that the process of lipid loading in Mφs is an intrinsically inflammatory process (3), yet recent studies have challenged this concept (4-7). Mechanistically, the metabolism of Mφs that accumulate excess intracellular cholesterol was shown to be rewired differently upon their activation. Specifically, their induction of AKT-dependent and HIF-1α-dependent glycolysis were significantly suppressed (5, 6). However, how they each contribute to the suppression of inflammatory responses in cholesterol loaded Mφs remains to be determined. In this study, we have dissected their relative contribution and found that while cholesterol loading of Mφs partially impaired to AKT-dependent glycolytic reprogramming, it is unlikely that it contributes to the subsequent suppression of HIF-1α transcriptional response. This is supported by the fact that complete inhibition of AKT only resulted in modest suppression of LPS-induced HIF-1α levels, glycolysis and pro-inflammatory gene expression. Moreover, the rate of synthesis of HIF-1α levels was not altered by cholesterol loading, yet its degradation was highly enhanced. Similarly, HIF-1α mutation studies also support that blocking its degradation is key to restoring HIF-1α levels in cholesterol loaded Mφs. Finally, our study also provided further evidence that cholesterol loading of Mφs alone is sufficient to induce oxidative stress and lipid peroxidation products, such as 4-HNE, and this is associated with a decline of reduced KEAP1 protein levels.

AKT has been proposed as a Warburg kinase due to its multi-faceted role in regulating cellular metabolism, such as rapidly enhancing the glycolytic flux through its kinase activity (24). Specifically, AKT has been shown to phosphorylate a variety of glycolytic enzymes, such as hexokinases (25), glucose transporters (26), phosphofructokinase (27). Apart from this, the hyperactivity of AKT has also been shown to regulate mTORC1 activity, which can directly increase its protein synthesis of HIF-1α through regulating the translation of its mRNA as it contains a 5’ TOP signal (13). However, the potential role of AKT in regulating HIF-1α abundance in the context of Mφ inflammation has not been explored and our study attempts to explore the causal relationship between these two events. To address this, we have utilized a pan-AKT inhibitor (MK2206) at a concentration that completely inhibited AKT activity and yet it only modestly inhibited HIF-1α protein levels. Understanding that the suppressive effect of cholesterol loading on AKT activation is far less potent than MK2206, it implicates the unlikelihood that the defective AKT signaling in cholesterol loaded PMφs directly contributes to the subsequent suppression of HIF-1α responses. Apart from this, we have also measured the synthesis rate of HIF-1α protein levels by inhibiting its proteasomal degradation in a time-dependent manner and did not find cholesterol loading modulated HIF-1α accumulation. This again not only reinforced the concept that the defective AKT signaling plays a minor role in regulating HIF-1α expression, but also demonstrated that other mechanisms that regulate HIF-1α synthesis, such as NF-кB-dependent transcription of HIF-1α mRNA (28), are also unlikely to be involved in regulating HIF-1α expression in cholesterol loaded Mφs. Our past research has shown that cholesterol loading of Mφs alone was sufficient to induce NRF2-dependent antioxidative stress response, and this consequently underlie the suppression of LPS-induced HIF-1α transcriptional responses (6). We further demonstrated that KEAP1, specifically its cysteine residue, such as C151, was dysfunctional as a result of cholesterol loading prior to LPS stimulation (6). However, it remains unclear how cholesterol loading-induced oxidative stress interferes with KEAP1 cysteine residues, and the molecular events that consequently occur after. In this study, we reasoned that lipid peroxidation species may play a role in interfering KEAP1 cysteine residues as the accumulation of excess cholesterol is often linked to the production of lipid peroxidation species (29). Among all, the effect of 4-HNE on inhibiting the function of KEAP1 is most established (30) and thus we measured the levels of proteins with 4-HNE adducts (formed via Michael addition or Schiff’s base formation) and found it was increased in cholesterol loaded PMφs. Based on past studies, 4-HNE has been shown to modify a variety of cysteine residues of KEAP1, such as C151 (31), a critical residue that we have previously shown to be affected by cholesterol loading, as well as C288 and C155 (32). Since our past study demonstrated that mutating C151 alone only partially blocked cholesterol loading-induced NRF2 response (6), it suggests that other cysteine residues of KEAP1 are likely to be involved and the broad reactivity of 4-HNE may contribute to the dysfunction of multiple cysteine residues. In addition to this, we have utilized redox western blotting to characterize the oxidized and reduced forms of KEAP1, and thus allow us to assess KEAP1 function during cholesterol loading. As shown in Figure 3E, cholesterol loading significantly increased oxidized KEAP1 proteins, while it nearly depleted all reduced KEAP1 proteins by the end of the cholesterol loading kinetic. This is strikingly different from the effects of H_2_O_2_-induced oxidative stress observed in Figure 3D, where the levels of oxidized and reduced KEAP1 protein levels returned as time passed and redox homeostasis was restored. In the cholesterol loaded PMφs, neither the oxidized nor the reduced forms of KEAP1 returned back to baseline, thus suggests the existence of a chronic oxidative stressed intracellular environment, where redox homeostasis was never achieved despite the induction of NRF2 response during cholesterol loading. Two possible reasons may explain the observations, one of which is the constant production of reactive oxygen species (ROS) during the course of lipid loading, which continuously react with multiple cysteine residues of KEAP1. This is supported by the findings from our past study that showed cholesterol loading-induced mitochondrial-derived ROS was still elevated by the end of the lipid loading kinetic (24h) (6). Another explanation is the formation of non-removable adducts on the cysteine residues of KEAP1, such as the ones formed by 4-HNE. 4-HNE is a powerful lipid peroxidation species partially due to its irreversible reaction, especially when 4-HNE is present at high concentrations (33). Therefore, this supports the possibility that cholesterol loading-induced 4-HNE irreversibly reacts with multiple cysteine residues of KEAP1, permanently locking it into an oxidized state and prevent it from being reduced, eventually leading to the observed depletion of reduced KEAP1 proteins at the end of the cholesterol loading kinetics. Future studies that utilize mass spectroscopy should be warranted to characterize the lipid peroxidation species and how they contribute to the post-translational modifications of KEAP1 cysteines induced by cholesterol loading of Mφs.

## Supplementary Figure Legends

**Supplementary Figure 1. The effect of AKT2 inhibition on LPS-induced pro-inflammatory and glycolysis genes. (A and B)** qPCR analysis of **(A)** pro-inflammatory (n=3) and **(B)** glycolysis gene expression (n=6) in ±AKT2i treated PMφs post 6h of LPS stimulation. The mean ± SEM is plotted in all graphs. Significant differences were determined by unpaired Student’s *t*-test in **(A and B)** (* P < 0.05, *** P < 0.001).

## Author Contributions

Conceptualization, K.K.Y.T., M.I.C.; Methods, K.K.Y.T.; Investigation, K.K.Y.T.; Formal Analysis, K.K.Y.T.; Visualization, K.K.Y.T; Writing – Original Draft, K.K.Y.T.; Writing – Review & Editing, K.K.Y.T., M.I.C.; Supervision, M.I.C.; Funding Acquisition, M.I.C

## Data availability statement

All data generated or analyzed during this study are included in this published article (and its Supplementary Information files)

