## Supplementary figures and images for "Cholesterol accumulation-induced impairment of AKT signaling in LPS-stimulated macrophages play a dispensable role in suppressing HIF-1α-dependent glycolysis"

### Supplementary Figure 1

#### Supplementary Figure 1

**A**

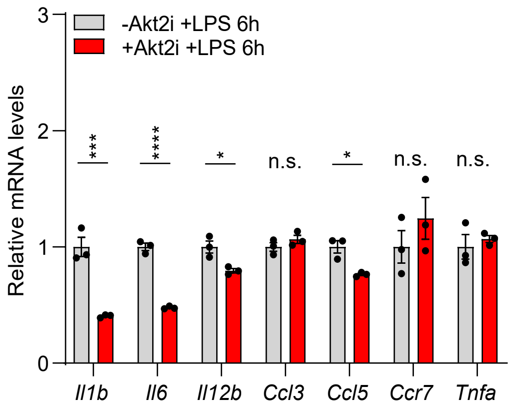


**B**

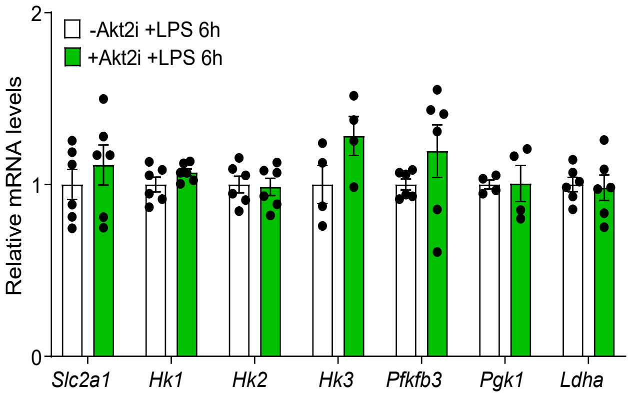
